# A unifying motif for spatial and directional surround suppression

**DOI:** 10.1101/180125

**Authors:** Liu D. Liu, Kenneth D. Miller, Christopher C. Pack

## Abstract

In the visual system, the response to a stimulus in a neuron’s receptive field can be modulated by stimulus context, and the strength of these contextual influences vary with stimulus intensity. Recent work has shown how a theoretical model, the stabilized supralinear network (SSN), can account for such modulatory influences, using a small set of computational mechanisms. While the predictions of the SSN have been confirmed in primary visual cortex (V1), its computational principles apply with equal validity to any cortical structure. We have therefore tested the generality of the SSN by examining modulatory influences in the middle temporal area (MT) of the macaque visual cortex, using electrophysiological recordings and pharmacological manipulations. We developed a novel stimulus that can be adjusted parametrically to be larger or smaller in the space of all possible motion directions. We found, as predicted by the SSN, that MT neurons integrate across motion directions for low-contrast stimuli, but that they exhibit suppression by the same stimuli when they are high in contrast. These results are analogous to those found in visual cortex when stimulus size is varied in the space domain. We further tested the mechanisms of inhibition using pharmacologically manipulations of inhibitory efficacy. As predicted by the SSN, local manipulation of inhibitory strength altered firing rates, but did not change the strength of surround suppression. These results are consistent with the idea that the SSN can account for modulatory influences along different stimulus dimensions and in different cortical areas.

**Significance Statement:** Visual neurons are selective for specific stimulus features in a region of visual space known as the receptive field, but can be modulated by stimuli outside of the receptive field. The SSN model has been proposed to account for these and other modulatory influences, and tested in V1. As this model is not specific to any particular stimulus feature or brain region, we wondered whether similar modulatory influences might be observed for other stimulus dimensions and other regions. We tested for specific patterns of modulatory influences in the domain of motion direction, using electrophysiological recordings from MT. Our data confirm the predictions of the SSN in MT, suggesting that the SSN computations might be a generic feature of sensory cortex.

## Introduction

What circuitry underlies sensory cortical processing? Recent work argues that visual cortical circuitry is well described by a circuit termed the Stabilized Supralinear Network (SSN) (Ahmadian et al., 2013; Rubin et al., 2015). The key idea is that neuronal gain – the change in output per change in input – increases with activation. As a result, the effective connection strengths between neurons increases with network activation, leading to a wide range of cortical nonlinear behaviors.

One such behavior involves surround suppression: a decrease in a neuron’s firing rate when the size of a stimulus exceeds that of the receptive field “center” (Allman et al., 1985; Jones et al., 2001; Cavanaugh et al., 2002). In the visual cortex, surround suppression is stronger for strong (high-contrast) stimuli than for weak (low-contrast) stimuli, so that the optimal stimulus size is larger for weaker stimuli (Sceniak et al., 1999; Pack et al., 2005; Tsui and Pack, 2011).

The SSN circuit explains this observation as follows. For very weak center stimuli, the cortical region representing the center is weakly activated and has weak effective connection strengths. Therefore, monosynaptic inputs to the center from the surround, which are primarily excitatory, dominate over di- and polysynaptic surround-driven local inputs, which are often inhibitory. As a result, the surround stimulus facilitates the response. With increasingly strong center activation, due either to a larger or higher-contrast stimulus, recurrent interactions become increasingly strong and increasingly inhibition-dominated (as observed in mouse V1, Adesnik (2017)). The surround stimulus then more strongly drives inhibitory neurons, yielding surround suppression. Thus, contrast-dependent surround suppression emerges from the dynamics of recurrent activity, without the need for explicit assumptions about different contrast thresholds for excitation and inhibition (Rubin et al., 2015).

Although the model has been primarily tested with V1 data, the underlying principles are generic (Ozeki et al., 2009; Rubin et al., 2015; Miller, 2016). In particular, if the connection strength between neurons decreases with their distance in a feature space (e.g., preferred orientation in V1, (Cossell et al., 2015); or preferred direction in MT), then the SSN model predicts that there should be contrast-dependent surround suppression in that feature space, just as in retinotopic space (Rubin et al., 2015). MT should show such a decrease in connection strength with increasing difference in preferred direction, because MT contains a local columnar structure (Albright, 1984) so that nearby neurons encode similar motion directions (Born and Bradley, 2005). The SSN thus predicts that MT neurons should show contrast-dependent surround suppression in the space of motion-direction: stimuli that include a wider range of motion directions, and thus activate MT neurons with a wider range of motion preferences, should suppress MT responses; and this direction-domain suppression should be stronger at higher contrasts and become weaker or absent at lower contrasts. Here we test this prediction in monkey area MT.

We also test a second prediction. For reasonably strong activation, the excitatory recurrence becomes strong enough that the network becomes an inhibition-stabilized network (ISN): a network in which recurrent excitation is strong enough to be unstable (i.e., epileptic), but the network is stabilized by feedback inhibition (Tsodyks et al., 1997; Ozeki et al., 2009). An ISN shows a “paradoxical” response: when external excitatory drive is added to inhibitory cells (as when a surround stimulus drives center inhibitory cells sufficiently strongly to cause surround suppression), the inhibitory cells *lower* their sustained firing rates, due to loss of recurrent excitation from suppressed excitatory cells. Thus, both excitatory and inhibitory cells are surround suppressed, as assayed by the inhibition received by excitatory cells being reduced by surround suppression (Ozeki et al., 2009; Adesnik, 2017). The SSN, and any model that is an ISN, predicts that surround suppression is little affected by *local* blockade of GABAergic inputs (Ozeki et al., 2004; Ozeki et al., 2009; Rubin et al., 2015), because the suppression is caused by a withdrawal of excitatory input that is not disrupted by local manipulations of inhibition.

We tested the first prediction by designing a stimulus that could be manipulated parametrically to be larger or smaller in the space of directions, while maintaining a fixed size in visual space. We found that responses in MT were indeed suppressed by stimuli with a wider range of motion directions, but only when the stimulus was high in contrast. At low contrast, neurons integrated over a larger spread of motion directions, as has been observed for spatial integration (Levitt and Lund, 1997; Kapadia et al., 1999; Sceniak et al., 1999). In addition, we confirmed that local blockade of GABAergic inhibition does not reduce spatial surround suppression in MT, just as in V1 (Ozeki et al., 2004). These results are consistent with the idea that the SSN is a generic mechanism of cortical computation (Miller, 2016).

## Materials and Methods

### Electrophysiological Recordings and Visual Stimuli

Two adult female rhesus monkeys (*Macaca mulatta,* both 7 kg) were used for electrophysiological recordings in this study. Before training, under general anesthesia, an MRI-compatible titanium head post was attached to each monkey’s skull. The head posts served to stabilize their heads during subsequent training and experimental sessions. For both monkeys, eye movements were monitored with an EyeLink1000 infrared eye tracking system (SR Research) with a sampling rate of 1,000 Hz. All procedures conformed to regulations established by the Canadian Council on Animal Care and were approved by the Institutional Animal Care Committee of the Montreal Neurological Institute.

Area MT was identified based on an anatomical MRI scan, as well as depth, prevalence of direction-selective neurons, receptive field size to eccentricity relationship, and white matter to grey matter transition from a dorsal-posterior approach. We recorded single units using linear microelectrode arrays (V-Probe, Plexon) with 16 contacts.

Neural signals were thresholded online, and spikes were assigned to single units by a template-matching algorithm (Plexon MAP System). Offline, spikes were manually sorted using a combination of automated template matching, visual inspection of waveform, clustering in the space defined by the principle components, and absolute refractory period (1 ms) violations (Plexon Offline Sorter).

Visual motion stimuli were displayed at 60 Hz at a resolution of 1,280 by 800 pixels; the viewing area subtended 60° × 40° at a viewing distance of 50 cm. Stimuli consisted of random dot stimuli displayed on a gray background (luminance of 98.8 cd/m^2^). Half the dots were black, and half the dots were white, resulting in a constant mean luminance across stimulus conditions. At 100% contrast, the black dots had luminance of 0.4 cd/m^2^, and the white dots had luminance of 198 cd/m^2^. The intermediate contrasts were defined as a percentage of the luminance difference from the gray background luminance, contrast = |(luminance - 98.8 cd/m^2^) / 98.8 cd/m^2^|. Animals were trained to fixate on a small dot at the center of the screen. Stimuli were shown after 300 ms of fixation. Each stimulus was presented for 500 ms, and the animals were required to maintain fixation throughout the stimulus and for another 300 ms after the end of the stimulus to receive a liquid reward. In all trials, gaze was required to remain within 2° of the fixation point in order for the reward to be dispensed. Data from trials with broken fixation were discarded.

The direction tuning and contrast response of the single units were quantified using 100% coherent dot patches placed inside the receptive fields. Offline the receptive field locations were further quantified by fitting a spatial Gaussian to the neuronal response measured over a 5 x 5 grid of stimulus positions. The grid consisted of moving dot patches centered on the initially hand-mapped receptive field locations. We confirmed that all neurons included in our analysis had receptive field centers within the stimulus patch used.

### Size Tuning Stimuli in Direction Space

We designed a stimulus that would allow us to study surround suppression in the motion domain in a manner that was analogous to studies in the spatial domain. In this conception, the input to the receptive field “center” is the strength of motion in a range about the neuron’s preferred direction. The “surround” is then motion in other directions, and the bandwidth of the center plus surround is the size of the stimulus in direction space. That is, a stimulus that contains motion in a range of directions spanning 180° is larger than a stimulus that spans a range of 60°. For these experiments we did not manipulate the spatial size of the stimulus, but rather fixed it according to the size of the hand-mapped spatial receptive field.

Our stimuli made use of random dots, each of which could be assigned to either a noise or a signal pool. The noise dots moved in random directions. The signal dots moved in a range of directions that straddled the preferred direction of each neuron. All dots moved at the same fixed speed of 8 or 16°/s, depending on the speed preference of the neuron. In all cases, dot patches were centered on the receptive fields determined by hand mapping. All conditions were interleaved randomly, and each stimulus was repeated 20 times.

We wished to change the size of the stimulus in direction space without changing other stimulus variables to which the neurons were sensitive. However, changing the size in direction space entails changing other low-level stimulus parameters (e.g., total number of dots or total amount of motion energy), which could confound our interpretation of the data. We therefore used two different methods to vary the stimulus bandwidth in direction space, each of which entailed changing a different low-level aspect of the stimulus.

In the first method, we kept the total number of stimulus dots fixed, and increased the motion bandwidth by drawing dots from a noise pool. Thus the total number of dots was identical for all stimuli, across variations in direction bandwidth. We constructed stimuli that contained signal dots moving in 1, 3, 5, and 7 directions, and each increase in the number of motion directions involved recruiting 25% of the noise dots to move coherently in the new direction (Fig. 1A and Table 1). This paradigm thus allowed us to test the influence of size in direction space for stimuli comprised of a fixed number of dots and a fixed amount of overall motion energy. We limited the largest size in direction space to be ±90° from the preferreddirection in order to avoid null direction suppression at larger sizes (Snowden et al., 1991; Qian and Andersen, 1994).

**Figure 1.**
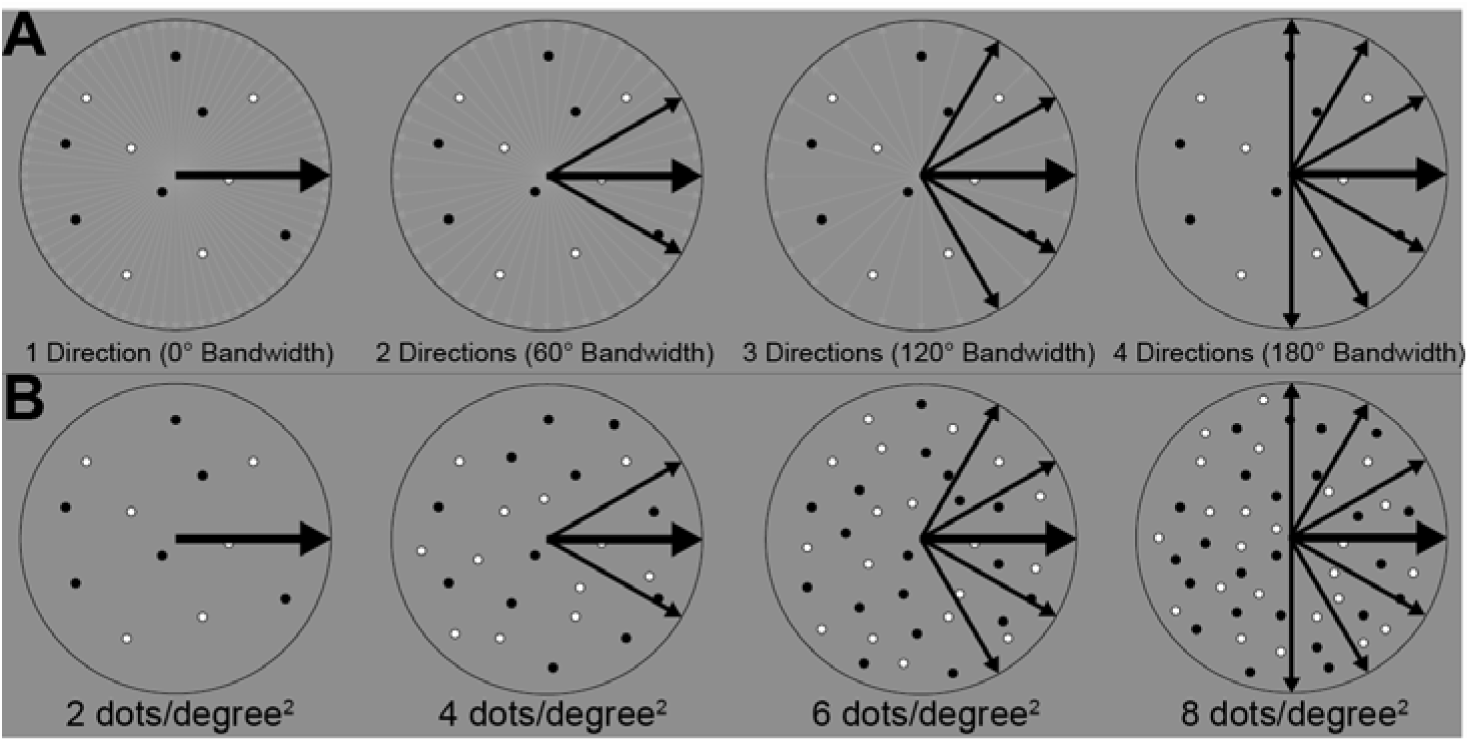
Illustration of the two methods of stimulus generation. **A,** Illustration of the stimulus that engages directional surround suppression in MT while the dot density is fixed. **B,** Illustration of the stimulus that engages directional surround suppression in MT while the dot density increases with directional size.

**Table 1.** Summary of the two methods of stimulus generation.

| Number of directions | Method 1: varying the noise pool, with dot density fixed to 2 dots/degree<sup>2</sup> (Fig 1A) |  | Method 2: varying the dot density without adding any noise dots (Fig 1B) |  |
| --- | --- | --- | --- | --- |
|  | Signal directions | Noise | Directions | Density |
| 1 | 25% at preferred direction | 75% | Preferred direction | 2 dots/degree<sup>2</sup> |
| 3 | 25% at preferred; 25% at ±30° from preferred | 50% | Preferred; ±30° from preferred | 4 dots/degree<sup>2</sup> |
| 5 | 25% at preferred; 25% at $\pm 30^\circ$ and 25% at $\pm 60^\circ$ from preferred | 25% | Preferred; $\pm 30^\circ$ and $\pm 60^\circ$ from preferred | 6 dots/degree<sup>2</sup> |
| 7 | 25% at preferred; 25% at $\pm 30^\circ$; 25% at $\pm 60^\circ$; and 25% at $\pm 90^\circ$ from preferred | 0% | Preferred; $\pm 30^\circ$, $\pm 60^\circ$ and $\pm 90^\circ$ from preferred | 8 dots/degree<sup>2</sup> |

However, in this approach, increases in motion bandwidth are yoked to decreases in noise, which might be expected to affect the strength of inhibitory inputs on their own (Hunter and Born, 2011). Thus, we also tested neurons using a second method, in which there was no noise pool, and we increased the size in direction space by simply adding more dots that moved in different directions. In this case the center stimulus strength (i.e. the strength of motion in the preferred direction) was constant across conditions, but the total number of dots (and hence the total motion energy) increased with stimulus size. The lowest dot density used was 2 dots/degree^2^, which is beyond the density at which MT responses typically saturate, at least for 100% coherence stimuli (Snowden et al., 1992). We again tested four different direction conditions (Fig. 1B and Table 1). In all cases, the dot size was 0.1°. The dots were initially plotted at random locations and moved in fixed directions from frame to frame. A dot that left the patch was replotted at the corresponding location on the opposite boundary of the patch on the next frame and continued its motion from there, i.e. the lifetime was equal to the stimulus duration (Qian and Andersen, 1994).

For all size tuning experiments in direction space, we tested each of the 4 sizes at high and low contrasts. High contrast was defined as 100% contrast, and the low contrast was chosen online to be around the *c_50_* of the contrast response function obtained with the 100% coherent dot patch. Offline, we eliminated one neuron for which the response at the lowest contrast was below 2 standard deviations of the spontaneous baseline firing rate.

### Grating, plaid, and pattern selectivity

We tested a subset of MT neurons (n = 65) with a standard measure of motion integration, the plaid stimulus (Movshon et al., 1985). Direction selectivity for each neuron was first measured with a 100% contrast drifting sinusoidal grating of spatial frequency of 0.5 cycles/°. Stimulus size and temporal frequency were matched to the neuron’s preferences. Plaid stimuli were constructed by superimposing two gratings (Fig. 5A).

We used the standard approach to quantify the component and pattern selectivity of each neuron (Smith et al., 2005). The partial correlations for the pattern and component predictions were calculated as,

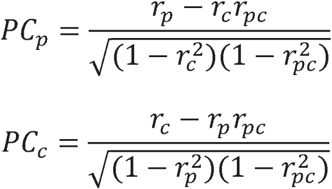

Here, *r_p_* and *r_c_* are the correlations between the plaid response and the pattern and component predictions, respectively, and *r_pc_* is the correlation between the pattern and component predictions. The partial correlations are z-scored as,

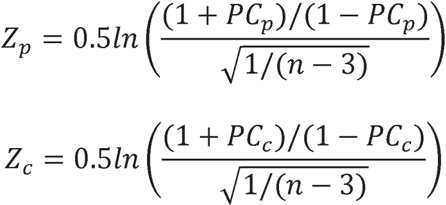

Where *n* = 12 is the number of directions. The pattern index was calculated as *Z_p_* — *Z_c_*.

### Pharmacological Injections

The pharmacological injection system has been previously described (Liu and Pack, 2017). Briefly, our linear electrode arrays contained a glass capillary with an inner diameter of 40 μm. One end of the capillary was positioned at the opening between contacts 5 and 6 of the array (contact 1 was most dorsal-posterior), so that the separation of the injection site from the recording contacts ranged between 0 and 1000 μm. The other end of the capillary was connected via plastic tubing to a Hamilton syringe for the injection of pharmacological agents with a minipump.

To effectively manipulate neuronal responses without compromising isolation, we typically used injections of 0.1-0.2 μL at 0.05 μL/min. For GABA, we used a concentration of 25 mM, which reduced neural activity without silencing it completely (Bolz and Gilbert, 1986; Nealey and Maunsell, 1994). For gabazine, the concentration was 0.05 mM, and we used injections of approximately 0.5 μL at 0.05 μL/min. In a few cases, this induced unstable and synchronized responses in the nearby neurons (Chagnac-Amitai and Connors, 1989). The electrophysiological recordings in those sessions were not further analyzed here.

### Data Analysis

MT direction tuning curves *r(xd)* were characterized by fitting a Gaussian function to the mean responses using the least-squares minimization algorithm (lsqcurvefit in MATLAB). The Gaussian function is

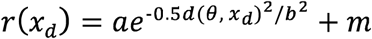

where *a* scales the height of the tuning curve; *b* determines the tuning curve width, the direction tuning width (DW) was defined as full width at half maximum of the fit, i.e. 2.35*b*; *x_d_* is the motion direction; *θ* is the preferred direction of motion; and *m* is the baseline firing rate of the cell. d(*θ, x_d_*) is the shortest distance around the 360 degree circle between *θ* and *x_d_*. The Gaussian fit to the data was very good in most cases (Median R^2^ = 0.90 before gabazine injection and R^2^ = 0.89 after injection).

The contrast response functions *r(x_c_)* were fitted with a Naka-Rushton function,

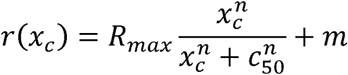

where *R_max_* scales the height of the contrast response function; *n* determines the slope; *c_50_* is the contrast at which the response function achieves half of its maximum response; and *m* is the baseline firing rate of the cell. *x_c_* is the contrast.

The neuronal size tuning curves *r(x_s_)* in retinotopic space were fitted by a Difference of Error functions (DoE) (Sceniak et al., 1999; DeAngelis and Uka, 2003),

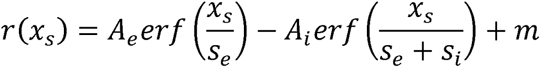

where *A_e_* and *A_i_* scale the height of the excitatory center and inhibitory surround, respectively. *s_e_* and *s_i_* are the excitatory and inhibitory sizes, and *m* is the baseline firing rate of the cell. *x_s_* is the stimulus size. The DoE fit to the data was very good in most cases (Median R^2^ = 0.93 before gabazine injection and R^2^ = 0.93 after injection).

The size suppression index (SI_S_) for each neuronal size tuning curve was calculated as SI_S_ = (R_m_ – R_L_)/R_m_, where R_m_ is the maximum across responses to different stimulus sizes and R_L_ is the response observed at the largest size. Since using the raw responses is sensitive to noise at both the maximum response and the response at the largest size, we used the values from the DoE fits for SI calculations.

Since we only measured the response at 4 sizes in the directional space, we were unable to fit a DoE function to the directional size tuning curves. Instead, to capture potential suppressive influences in the direction domain, we calculated a direction integration index from the raw data II_D_ = (R_L_ – R_S_) / (R_L_ + R_S_), where R_L_ is the response observed at the largest size and R_S_ is the response observed at the smallest size.

### SSN Model Simulations

We first simulated a 1D ring model, which captures putative interactions among neurons representing different motion directions (Fig. 2A). Details of this model can be found elsewhere (Rubin et al., 2015). Our model differs in that the ring is 360 degrees in extent (vs. 180 degrees in Rubin et al., 2015), representing all possible motion directions. There is an excitatory (E) and inhibitory (I) neuron at every integer position *x_i_* = 0°, 1°, …, 359°, where *x_i_* represents the preferred direction of the corresponding E and I cells. We can write the model equation in matrix notation as,

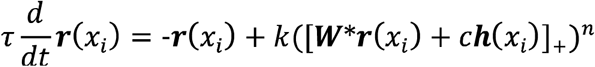

where ***r***(*x_i_*) is the vector of firing rates of the excitatory and inhibitory neurons with preferred motion direction *x_i_, **W***(*y*) is the weight matrix of E → E, E → I, I → E, and I → I connections between neurons separated by angular distance y (measured as shortest distance around the 360° circle). The connection weights W_ab_(y) = J_ab_Gσ_dir_(y), where J_EE_ = 0.044, J_EI_ = 0.023, J_IE_ = 0.042, J_II_ = 0.018, Gσdir(y) are a Gaussian function with standard deviation of 64° (Ahmadian et al., 2013). ***W*r***(*x_i_*) is the convolution **Σ_*j*_*W***(*x_i_*-*x_j_*)**r**(*x_j_*) where the sum is over all preferred directions *x_j_*; ***h***(*xi*) is the vector of external input to the E and I neurons preferring *x_i_*; and *c* is the strength (monotonically related to contrast) of the input. The elements of the vector of input to the neuron, ***W*r***(*x_i_*) + ***ch***(*x_i_*), are thresholded at zero before being raised to the power *n*: [z]_+_ = 0 *if z* < 0, = *z if z* ≥ 0 (the operations of thresholding and raising to a power are applied separately to each element of the vector). *k* and *n* are identical for E and I neurons, with *k* = 0.04 and *n* = 2. *τ* is a diagonal matrix of the time constant for E cells, τ_E_ = 20 ms, and for I cells, τ_I_ = 10 ms.

**Figure 2.**
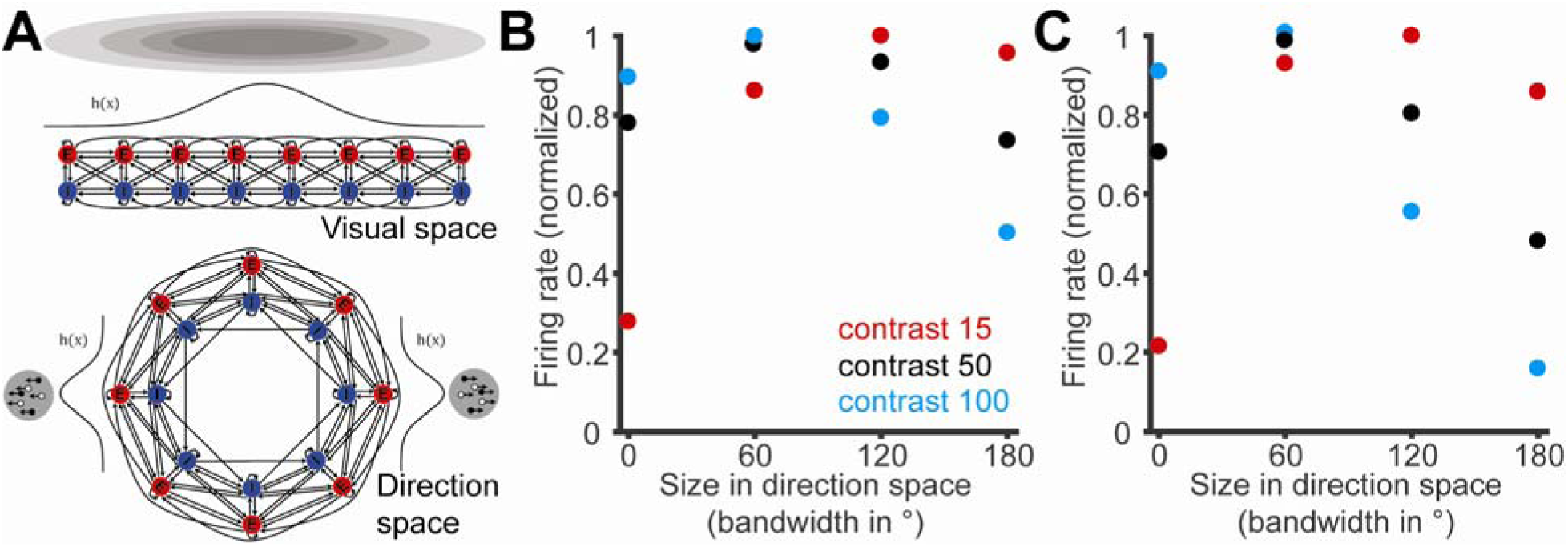
Stabilized supralinear network can account for surround suppression in both spatial and direction domains. **A,** Schematic of the 1D SSN ring model as a direction space analogue of the visual space model. In the visual space model (top), stimuli of different sizes in visual space (gray circles) are simulated as input, *h(x),* of varying width, to a linear 1D grid of excitatory (E, red) and inhibitory (I, blue) units. The grid positions represent visual space positions. In the direction space (bottom), there are 360 E and I units, with coordinates on the ring as preferred directions. A dot stimulus, *h(x),* moving at a single direction is a Gaussian-shaped input with standard deviation of 60°. Stimuli including multiple directions simply add such input for each direction. We considered two methods of adding directions: including a “noise pool” stimulus of equal input to all directions, and subtracting from the noise pool as we added directions to keep total input strength unchanged (Fig. 1A); or simply adding additional input as we added directions, without a noise pool (Fig. 1B). **B,** Directional surround suppression at high contrast, but not at low contrast, arises from the dynamics of the model. This simulation result is for the first method of taking dots from a noise pool to add further directions about the preferred (Fig. 1A). The response at each contrast is normalized to the peak response. **C,** The simulation result for the second method of adding dots to further directions about the preferred without a noise pool (Fig. 1B). The response at each contrast is normalized to the peak response.

Regarding the model parameter choices, the four amplitudes J_ab_ were constrained to ensure stability and strong nonlinear behavior. To ensure stability, we require J_EI_J_IE_ > J_EE_J_II_, meaning feedback inhibition is sufficiently strong. For equal-strength inputs to E and I cells as used here, the strongest nonlinear behavior also requires J_II_ – J_EI_ < 0 and J_II_ – J_EI_ < J_IE_ – J_EE_ (Ahmadian et al., 2013). We chose Gσ_dir_(y) to have a standard deviation of 64°, given the bandwidth of MT direction tuning curves and the idea that cells with more strongly overlapping tuning curves should more strongly connect to each other; this value can be varied to give a diversity of surround suppression as observed in the data. We chose n = 2 for the power-law input-output (I/O) function, consistent with the observation in V1 that neurons have I/O functions well described by a power law throughout the full range of firing induced by visual stimuli, with powers in the range 2-5 (Priebe and Ferster, 2008). At n = 2, k =0.04 gave reasonable firing rates, but the qualitative behavior is consistent for a large range of n and k. Finally, we chose the ratio of the time constants for E and for I cells, τ_E/_τ_I_ = 2, to help ensure stability; given that the network is stable, the time constants do not affect the steady-state network responses, which is what we are modeling here.

We simulated network responses to random dot field stimuli of variable coherence. We assumed that a coherent dot stimulus of a given direction gives input to MT neurons proportional to a Gaussian function, of standard deviation 60°, of the difference (shortest distance around a 360° circle) between the neuron’s preferred direction and the stimulus direction. To simulate the method using noise dots (Table 1, Method 1), the non-coherent (noise) dots gave equal input, proportional to 1/360, to neurons of all preferred directions. The strength of the stimulus is given by a parameter *c*, identified as the “contrast” in Figure 2. As in our electrophysiological experiments, we used stimuli corresponding to 4 different sizes in direction space (Fig. 1A). Thus for the smallest size, 25% of the input, ***h***, was modelled as a Gaussian distribution around the preferred direction (peak of the Gaussian = *c*/4), while the remaining 75% was spread equally around the ring (uniform distribution of size (3/4) x *c*/360). At 2 directions, an additional 25% was taken from the non-coherent input and added to Gaussian spreads about +/-30° from the preferred direction (these two Gaussians have peak = *c*/8; noise amplitude becomes (1/2) x c/360). 3 and 4 directions followed in a similar manner while the total input strength was kept constant across sizes. We also simulated Method 2 (Table 1), which used the same set of stimuli except without a noise background (so that the total input strength grew with increasing number of directions), and the results were qualitatively similar as presented in Results.

### Experimental design and statistical analysis

We used two female rhesus monkeys (*Macaca mulatta*) for electrophysiological recordings in this study; this is standard for electrophysiological studies involving monkeys. We used the Wilcoxon rank-sum test to evaluate the difference between the Integration Index at low and high contrast, and the difference between Direction Tuning Width and Suppression Index before and after injection of Gabazine. As the Direction Tuning Width and Suppression Index can be affected by the ability to sample the tuning curves, we performed a bootstrapping analysis to ensure the robustness of the summary statistics. For each cell, we randomly sampled (with replacement) 10 trials per direction or size to create a tuning curve and then fitted a circular Gaussian or DoE to the subsampled tuning curve to generate a new direction tuning width or suppression index. We generated 100 sample distributions and tested the effects of gabazine injections with a Wilcoxon signed-rank test. To evaluate the relationship between the Pattern Index and Direction Tuning Width and the Integration Index, we calculated Pearson correlation coefficients. All analyses made use of built-in MATLAB functions and custom scripts. The complete results of the statistical analyses for each experiment can be found in the corresponding Results section.

## Results

In this section, we first present simulation results for the SSN. We then test a crucial model prediction with neurophysiological recordings from MT neurons in awake and behaving macaques. The theoretical and empirical results show that surround suppression in the motion domain behaves similarly to surround suppression in the space domain, with integration at low contrasts switching to suppression at high contrasts (Figs. 3 and 4). We also find that pattern-selective cells (as assayed from plaid responses) show greater motion integration than component-selective cells (Fig. 5). Finally, as predicted by the SSN model, local pharmacological manipulation of inhibition does not alter spatial surround suppression, although our methods had the expected effects on directional tuning width (Figs. 6 and 7).

### Stabilized supralinear network predicts contrast-dependent surround suppression in the direction domain in MT

Previous instantiations of the SSN have considered a model in which connections are defined either across a retinotopic sheet of the kind found in V1 or across a ring of preferred orientations (Ahmadian et al., 2013; Rubin et al., 2015; Miller, 2016). Like orientation, motion direction is a circular variable, but it takes values over 360° rather than 180° as for orientation. Thus to examine the properties of the SSN in this circular space, we first simulated a ring model ((Rubin et al., 2015); Fig. 2A) of motion direction space. This represents neurons of varying preferred directions sharing a common location in retinotopic space.

In general, the SSN predicts that contrast-dependent surround suppression should occur in any stimulus feature dimension, provided certain minimal connectivity conditions are met, e.g. average connection strength between neurons decreases with the dimensional distance between them. We accordingly assumed that the strengths of connections between neurons on the ring decreased with increasing difference in their preferred directions. By analogy with the study of size-tuning in the spatial domain, we tested the SSN with stimuli of different motion-domain sizes. We increased the size of the stimulus in direction space by including stimuli at increasingly wider ranges of directions about the preferred direction (the “center” of the receptive field). As described in Methods, we considered size or bandwidth 0° (preferred-direction stimulus only), 60° (adding stimuli at +/− 30° about the preferred), 120° (adding additional stimuli at +/− 60°), and 180° (additional stimuli at +/-90°). For each motion size, we examined different levels of stimulus contrast, represented as scaling the strengths of all inputs.

The simulation results (Fig. 2B) show that the model predicts strong direction-domain surround suppression at high contrast, but not at low contrast. Specifically, at low contrasts (red), increasing the range of motion directions leads to increased responses with a hint of suppression for the largest stimulus size, while at high contrasts larger motion-domain stimulus sizes lead to strong suppression (blue). Intermediate contrasts give an intermediate result (black). These results change very little with changes in the total number of dots in the stimulus (Fig. 2C), a factor that we consider in our experiments below (Fig. 4). Thus the model consistently predicts direction-domain suppression that is analogous to space-domain surround suppression. In the SSN, the dependence of surround suppression on contrast arises generically from the dynamics of the SSN in summing inputs, rather than by the assumption of a higher contrast threshold for inhibition, as in previous models (Somers et al., 1998; Huang et al., 2008; Schwabe et al., 2010; Carandini and Heeger, 2012).

### Surround suppression in direction domain of MT

We tested the model predictions by recording from individual MT neurons, using the same stimuli as in the simulations. We first show results for the first type of stimulus described above, in which there was a noise pool of dots moving in random directions. For each neuron we fixed the physical size of each stimulus according to an estimate of the classical receptive field size. We then varied stimulus size in the motion domain, as well as dot contrast. Thus for the smallest stimulus, all the coherent dots moved in the preferred direction of the neuron (Fig. 1A, left), with the remaining dots in the noise pool moving in random directions. To increase the size of stimuli in the motion space, we recruited dots from the noise pool and added them to directions around the preferred direction (Fig. 1A). This manipulation kept the total motion energy and dot density of the stimulus constant across sizes.

Figure 3A shows the firing rate of an example MT neuron for stimuli of different contrasts and motion sizes. For the low contrast stimulus (red), firing rate increased with motion size, while for higher contrasts (blue, black) firing rate decreased with motion size. Thus the pattern of firing rates for this neuron was consistent with the SSN prediction that MT neurons would shift from motion-domain integration to suppression as the stimulus contrast was increased (Fig. 3A). Indeed, just as in the space domain, for large stimuli it is possible to increase firing rates by lowering contrast (Fig. 3A; Pack et al., 2005).

**Figure 3.**
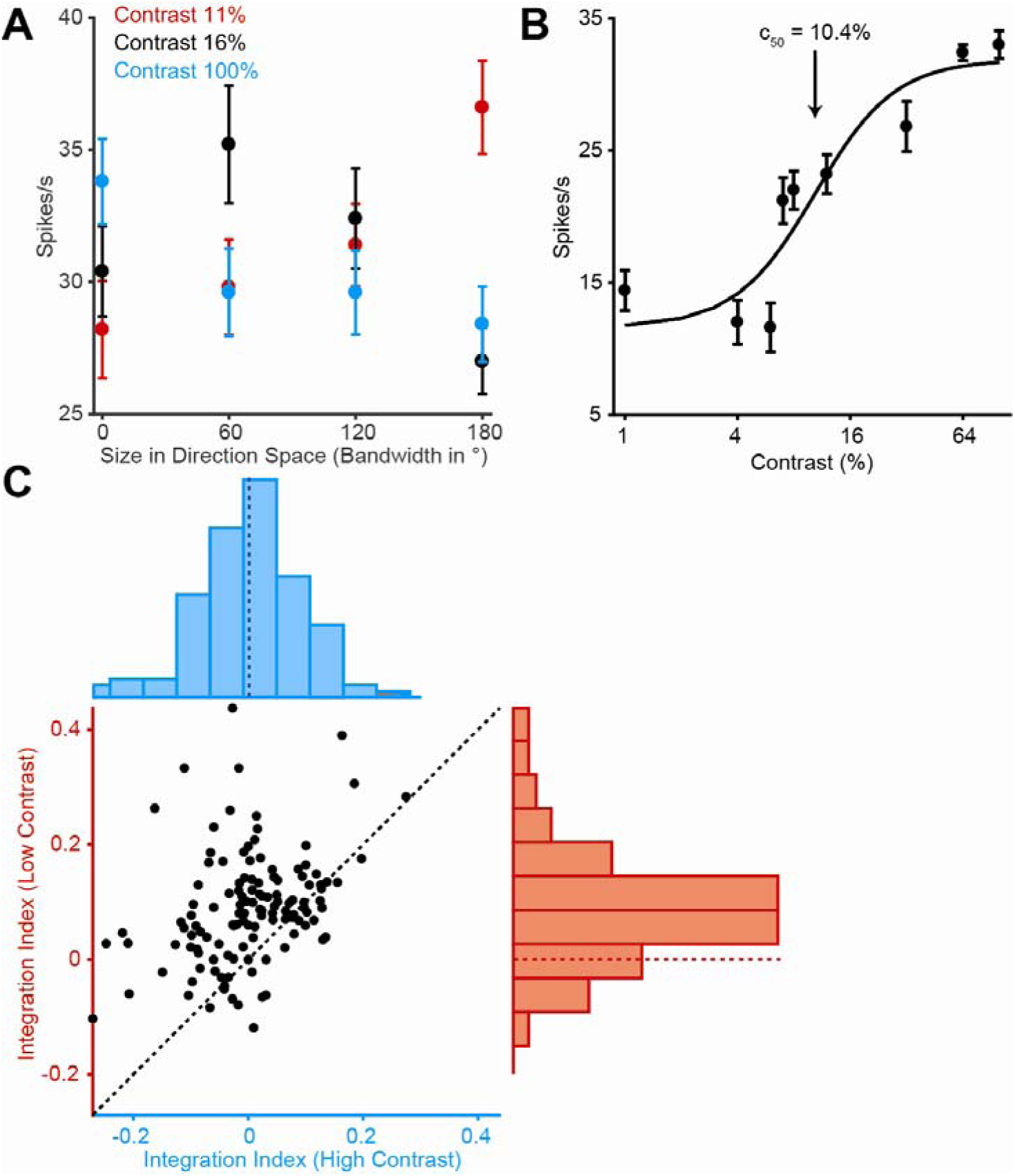
Surround integration and suppression in the direction domain. **A,** Surround suppression occurs in direction space at high contrast, but not at low contrast for an example neuron. **B,** Contrast response function for the same example neuron using 100% coherent dots in the preferred direction. The line indicates the Naka-Rushton function fit. **C,** Population data for direction surround integration. Scatter plot of the integration index, II_D_, at low contrast against the II_D_ at high contrast (rank sum test, *p* < 0.001). The marginal distributions are histograms of the IId (Median at high contrast = 0.002; Median at low contrast = 0.084). Dashed lines in the histograms show location of II_D_ = 0.

To examine these effects across the MT population, we calculated the directional integration index (II_D_, the difference between responses to the largest and smallest sizes divided by the sum of these responses; see Methods) for data of the kind shown in Figure 3A for 125 neurons. The II_D_ captures the integration of signals across motion directions, with larger II_D_ values indicating more integration. Across the population (Fig. 3C) the II_D_ was frequently below zero, indicating a suppression of the response when dots activated the directional surround. Overall the II_D_ was significantly decreased at high contrast compared to low contrast, consistent with reduced integration at high contrasts (*p* < 0.001, rank sum test; *p* < 0.001 for monkey 1 and *p* = 0.01 for monkey 2). Note that this is not due to a failure of the low contrast stimuli to elicit a response from the neurons, as all neurons except one showed responses to the lowest contrast tested that were significantly above baseline. The one neuron that failed to meet this criterion was eliminated from further analysis. Overall, these results are similar to previous results in the space domain in MT (Pack et al., 2005; Tsui and Pack, 2011). However, the mechanisms of spatial and directional integration for a given cell appeared to be independent, as there was no correlation between the degree of spatial surround suppression and directional surround suppression measured at high contrast in the same neurons (Pearson’s *r* = -0.06, *p* = 0.46, N = 124).

We also tested 46 neurons using a second stimulus in which there was no noise pool, and we increased the total number of stimulus dots with size in the direction domain (Fig. 1B). This stimulus was designed to control for a potential confound in the previous experiment, which kept the total number of dots constant across stimulus size. In the latter configuration, increases in direction-domain size were yoked to decreases in the number of noise dots, and because noise includes motion in all directions, this can be viewed as reduction in the strength of the directional surround, analogous to the far surround in retinal space (Angelucci and Bullier, 2003; Angelucci and Bressloff, 2006). The new stimulus was directly analogous to that typically used in size tuning experiments, in which the stimulus is simply expanded to probe the influence of the surround.

We tested this subpopulation of MT neurons with both stimuli, and the results are shown in Figures 4A and 4B. For the control stimulus, the II_D_ is still significantly higher at low contrast than at high contrast (Fig. 4A; *p* = 0.04, rank sum test). Thus integration across direction space was greater at low contrast, regardless of how size was manipulated. For these neurons, we also replicated the previous result using the stimulus with a constant total number of dots (Fig. 4B; *p* < 0.001, rank sum test). The contrast modulation of II_D_ was not significantly different for the two stimulus types (rank sum test, *p* = 0.45).

**Figure 4.**
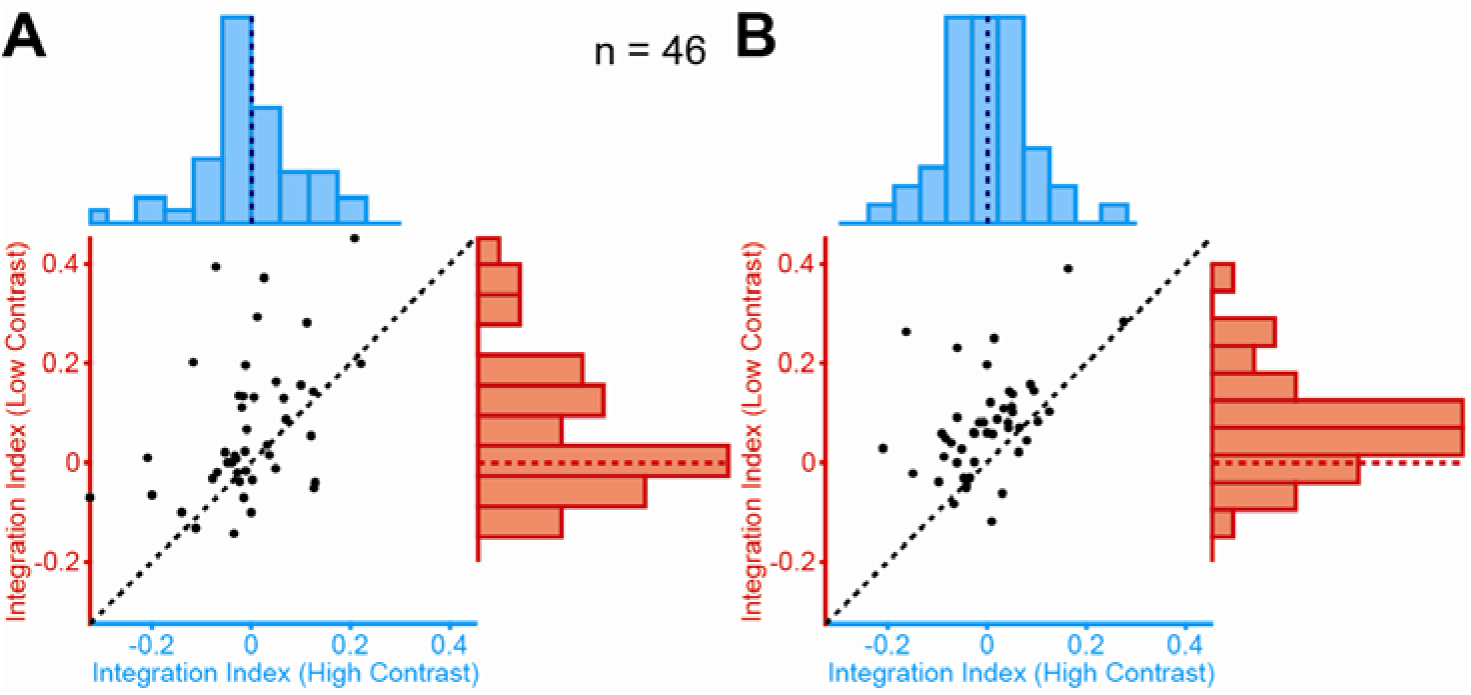
Additional controls for direction surround integration and suppression. **A,** Population data for direction surround integration. Scatter plot of the directional integration index (II_D_) at low contrast against the II_D_ at high contrast (rank sum test, *p* = 0.04). The marginal distributions are histograms of the II_D_ (Median at high contrast = -0.012; Median at low contrast = 0.018). Dashed lines in the histograms show location of II_D_ = 0. **B,** The contrast modulation of II_D_ for the same 46 neurons as in B, when the number of dots is held fixed by drawing from a noise pool (as in Fig. 3). The conventions are the same as in panel B (Median at high contrast = 0.003; Median at low contrast = 0.065).

Of the complete MT population, 65 were also tested with a standard probe of direction domain integration, the plaid stimulus (Movshon et al., 1985). Our plaid stimuli consisted of two superimposed sine-wave gratings, moving in directions separated by 120° (Fig. 5A); stimulus size was again matched to the classical receptive field, and contrast was 100%. From the resulting data we computed a pattern index (see Methods; Smith et al., 2005), which captures the extent to which MT neurons integrate the two motion directions; higher values indicate greater integration (Fig. 5B and C). We found that the pattern index was significantly correlated with the directional II_D_, as measured in our direction-size-tuning experiments at both low (Fig. 5D; Pearson’s *r* = 0.33, *p* = 0.01) and high contrasts (*r* = 0.27, *p* = 0.03). That is, cells with higher pattern indices showed less surround suppression in direction space – greater motion integration -- both at low and high stimulus contrasts. This suggests that area MT might use similar mechanisms to integrate motion signals for dot stimuli and grating stimuli. We also found that there was no correlation between the directional motion integration index and the width of the direction tuning curve, as measured using responses to standard stimuli of drifting dots moving coherently in a single direction (Fig. 5E; Pearson’s *r* = -0.08, *p* = 0.38 for low contrast, *r* = 0.05, *p* = 0.57 for high contrast).

**Figure 5.**
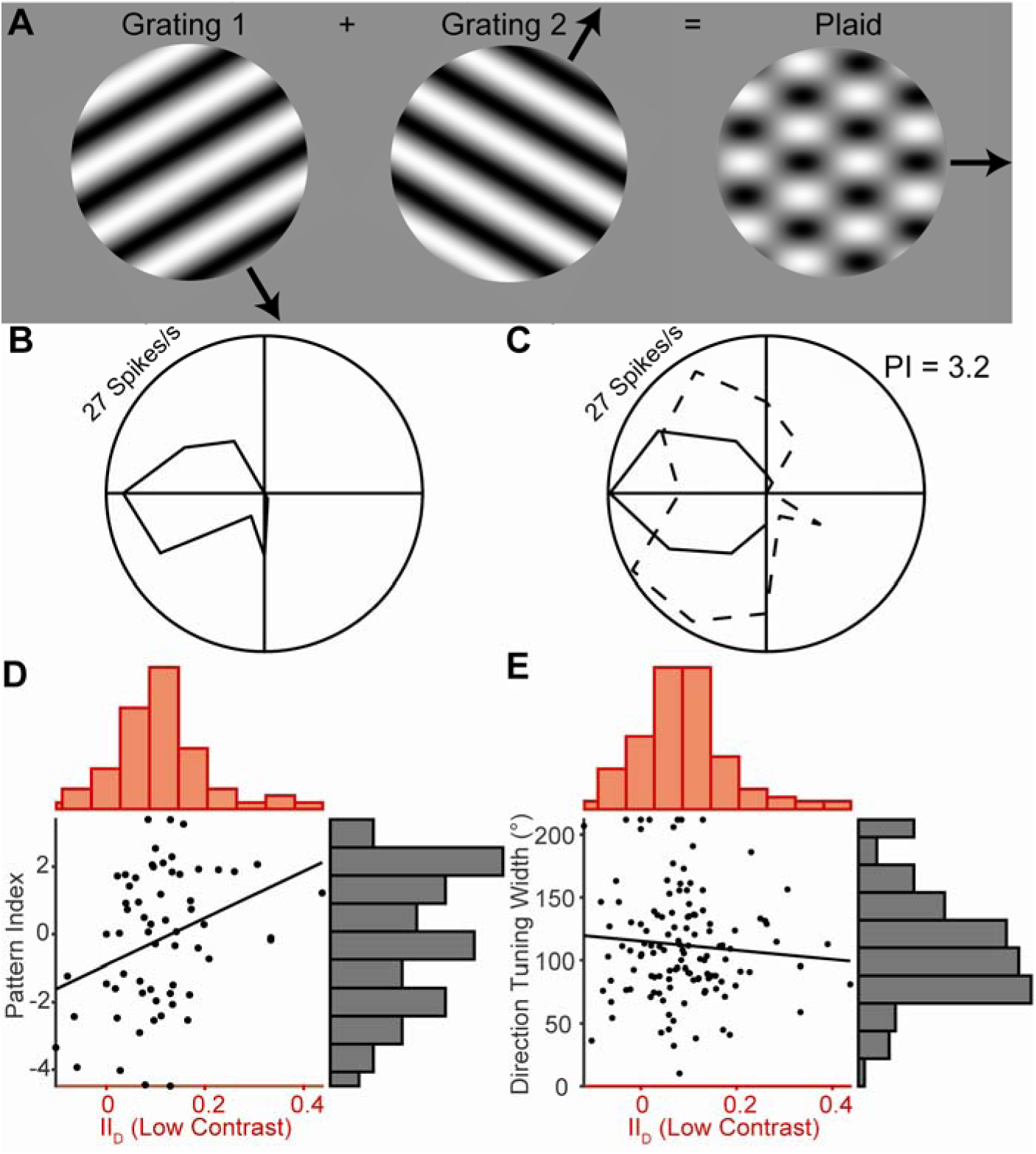
Direction integration with plaid stimuli. **A,** Illustration of the grating (left) and plaid stimuli (right). **B,** Direction tuning curve for an example neuron in response to drifting gratings. **C,** Direction tuning curve for the same neuron in response to moving plaids. The dashed line indicates the component prediction, which is the expected result if the neuron fails to integrate the motion of the plaid. **D,** Population data for motion integration. Scatter plot of the pattern index against the directional integration index (II_D_) at low contrast (*r* = 0.33, *p* = 0.01). **E**, Scatter plot of the direction tuning width against the directional integration index (II_D_) at low contrast (*r* = -0.08, *p* = 0.38).

### GABAergic influence on neuronal direction tuning and surround suppression in the spatial domain

Another prediction of the SSN is that local changes in the strength of inhibition should have little or no effect on surround suppression, because surround suppression is a result of withdrawal of network excitation (as well as inhibition), and a local blockade of inhibition will not change these network dynamics (Ozeki et al., 2009). This is different from conventional models, which posit that suppression is induced by an increase in the inhibition that a cell receives, so that a reduction in the inhibition to a given neuron will reduce its surround suppression (Tsui and Pack, 2011). Previous work has confirmed the SSN predictions in anesthetized cat V1, using iontophoretic injection of GABA antagonists: inhibitory blockade did not reduce surround suppression (Ozeki et al., 2004). In this section, we examine the effects of pharmacological manipulation of GABA in MT of awake monkeys.

We first confirmed that gabazine, a GABA_A_ receptor antagonist, robustly modulated neuronal firing in MT (Thiele et al., 2012). We measured direction tuning using random-dot stimuli of fixed spatial size, with all dots moving coherently in a single direction (Fig. 6A). We found that injection of gabazine increased direction tuning width, as found previously (Thiele et al., 2004; Thiele et al., 2012). In contrast, injections of GABA decreased firing rates across all directions (Fig. 6E), leading to narrower tuning (Leventhal et al., 2003).

**Figure 6.**
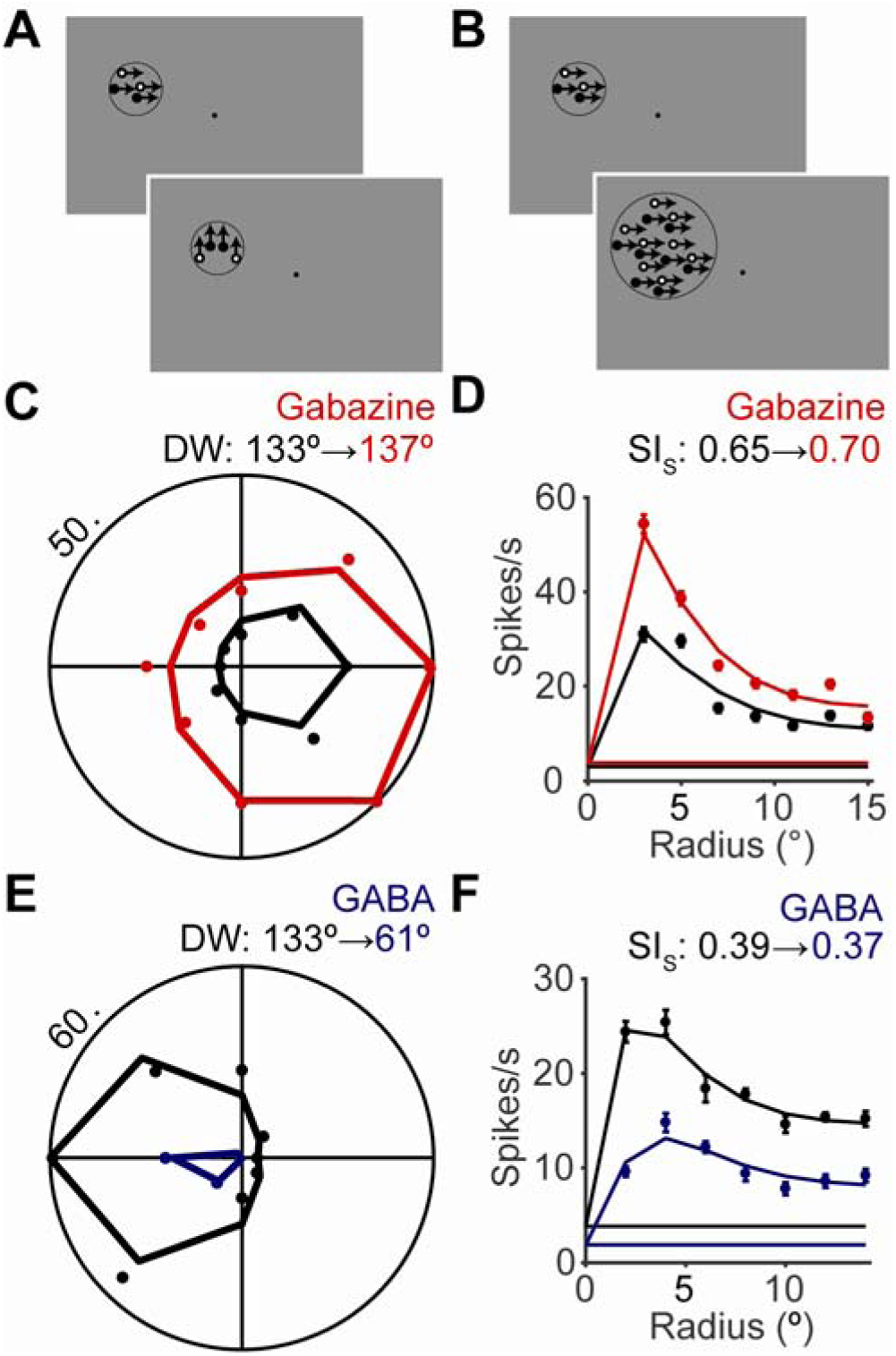
Effect of GABA on motion direction and size tuning. **A and B**, 100% coherent random dot patches were used to probe the direction and size tuning of MT neurons. **C and E**, Direction tuning curve for an example neuron before (black) and after injection of gabazine (C, red) or GABA (E, blue). The points are the mean responses for each direction. The lines indicate Gaussian function fits. Direction tuning width (DW) was defined as full width at half maximum of the fit. **D and F**, Size tuning curves for an example neuron, plotting the firing rate (mean ± s.e.m.) as a function of patch size before (black) and after injection of gabazine (D, red) or GABA (F, blue). The lines indicate difference of error functions fits. The horizontal lines show the spontaneous firing rate.

Figure 7A summarizes the influence of gabazine on direction tuning widths for a population of 38 MT cells: Tuning width increased following the injection, as determined by a rank sum test (*p* = 0.04) and verified with a bootstrapping analysis (see Methods; Wilcoxon signed-rank test; *p* < 0.001); these increases were particularly noticeable for cells that were narrowly tuned before the injection, as noted previously in V1 of anesthetized cat (Katzner et al., 2011). These changes in tuning width were not associated with changes in spontaneous firing rate, as the changes in spontaneous were modest and did not reach statistical significance (rank sum test, *p* = 0.32). Moreover, there was no correlation between gabazine-induced changes in spontaneous firing and changes in tuning width (Pearson’s *r* = 0.05, *p* = 0.78). We did not have enough data from the GABA experiments to perform statistical analyses, but in all 5 experiments, direction tuning width decreased following injection.

**Figure 7.**
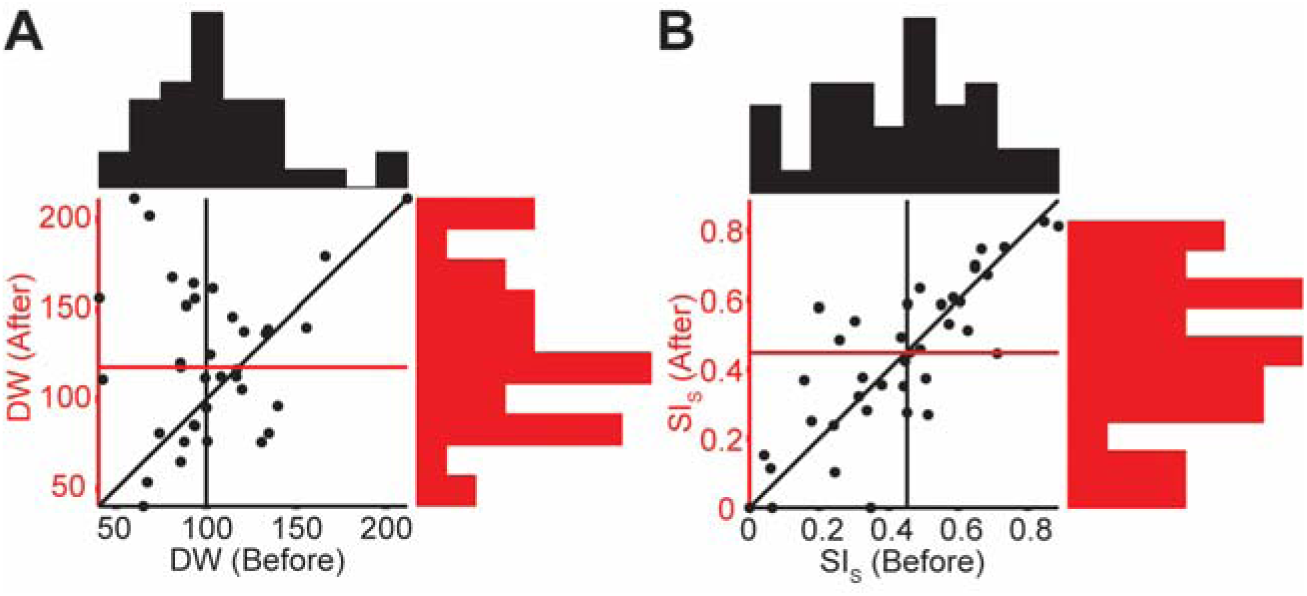
Population data on the effects of gabazine on direction and size tuning. **A**, Scatter plot of the direction tuning width before the injection of gabazine against the tuning width after injection (rank sum test, *p* = 0.04). Red and black lines represent the medians of the respective marginal distributions. **B,** Scatter plot of the neuronal size suppression index (SI_S_) before the injection of gabazine against the neuronal SI_S_ after injection (rank sum test, *p* = 0.98).

To test the influence of GABA concentrations on surround suppression, we performed standard (space-domain) measurements of size tuning, using random-dot stimuli (100% coherence) of different physical extents, with all dots moving in the neuron’s preferred direction (Fig. 6B). Previous work has shown that these stimuli elicit surround suppression in the upper and lower layers in MT, but not in layer 4, suggesting that the suppression is generated through intrinsic connections within MT (Born and Tootell, 1992; Raiguel et al., 1995). This property makes such stimuli useful for testing the predicted role of inhibitory inputs in the SSN.

Figure 6D shows size tuning curves from the same MT neuron as in Figure 6C. The preinjection data (black line) show that the neuron exhibited substantial surround suppression, as the response was reduced significantly with increasing stimulus size. As for the direction tuning curve, injection of gabazine increased firing rates in a non-specific manner. However, in this neuron there was no apparent reduction in surround suppression (Fig. 6D), and this result was generally true for the MT population (n = 38): The size suppression index (SI_S_), defined as the difference between the peak response and the response to the largest stimulus divided by the peak response, was similar before and after injection of gabazine (Fig. 7B; rank sum test, *p* = 0.98; bootstrapping analysis followed by Wilcoxon signed-rank test; *p* = 0.99). Again there was no correlation between the effects of gabazine on SI and the effects on spontaneous firing (Pearson’s *r* = -0.11, *p* = 0.52). These results are similar to those found in V1 of anesthetized cats (Ozeki et al., 2004), despite the much larger volume of gabazine used here. In a smaller sample (n = 5), we found that injection of GABA did not increase surround suppression, despite a strong overall reduction in firing rate (Fig. 6F).

## Discussion

Through electrophysiological recordings in awake monkeys, we have found contrast-dependent surround suppression in MT in a space defined by motion directions. In addition, we found that local manipulation of the efficacy of GABAergic inhibition had little influence on standard measures of surround suppression. Both results are consistent with predictions of the stabilized supralinear network (SSN), previously tested in V1 (Rubin et al., 2015).

### SSN as a unifying motif for normalization in multiple cortical areas

The contrast dependence of surround suppression in the space domain has been observed in both V1 and MT (Polat et al., 1998; Kapadia et al., 1999; Sceniak et al., 1999; Pack et al., 2005; Schwabe et al., 2010; Tsui and Pack, 2011). These results have previously been modeled under the assumption that inhibitory neurons have higher contrast thresholds than excitatory neurons (Somers et al., 1998; Huang et al., 2008; Schwabe et al., 2010; Carandini and Heeger, 2012). However, there is little experimental support for this assumption, and some data that contradict it (Contreras and Palmer, 2003; Song and Li, 2008).

In the SSN, the excitatory and inhibitory units can have the same properties (Rubin et al., 2015). Each unit has a power-law input/output function, but is stabilized by network inhibition (Ozeki et al., 2009; Ahmadian et al., 2013; Rubin et al., 2015). With low contrast inputs, the recurrent interactions within the network are weak, so neurons act relatively independently, summing their feedforward inputs and responding according to their transfer functions. With higher-contrast inputs, strong recurrent connections within the network provide contrast- and size-dependent suppression, with size in the spatial and feature (direction) domains playing similar roles.

The SSN also predicts that the local blockade of GABA_A_ receptors should not reduce surround suppression (Ozeki et al., 2009). In the SSN, surround suppression is not a result of an increase in inhibitory GABAergic input, but a withdrawal of both excitation and inhibition. In contrast, in models in which surround suppression results from an increase in the inhibition received by suppressed neurons (e.g., Tsui and Pack, 2011), local blockade of inhibition should reduce or prevent surround suppression.

Modulatory influences in visual cortex are often modeled within the normalization framework, which is hypothesized to be a generic computation with equal validity across brain regions and stimulus modalities (Carandini et al., 1997; Reynolds and Heeger, 2009; Carandini and Heeger, 2012; Krause and Pack, 2014). The normalization model as typically conceived, is a phenomenological rather than circuit model, in which some form of unnormalized neuronal response is suppressed by the sum of unnormalized responses in other neurons that constitute the “normalization pool”. The precise form of normalization, for example whether the normalizing pool constitutes all neurons or is restricted in some way based on neuronal tuning, must be matched to fit the particular experiments modeled.

The SSN can be regarded as a circuit instantiation of the normalization model, in that many SSN results closely match the results of an appropriately constructed normalization model (Rubin et al., 2015). In the circuit implementation, the form of normalization is determined by the connectivity. For example, in the SSN, orientation-specific long-range horizontal connectivity leads to the orientation-selectivity of surround suppression (Rubin et al., 2015); in a normalization model, this would be explained by assuming that the normalization pool consists of neurons of similar preferred orientations to the normalized cell. The normalization model does not explain the mechanism of suppression, and alternative mechanisms yield different predictions. For example, if the normalization pool exerted suppression by adding inhibition to the normalized cells, then one would expect increased inhibition and increased conductance in normalized (e.g., surround-suppressed) cells, and local GABAergic blockade would reduce or eliminate the normalization. In the SSN mechanism, normalization typically results from a decrease in both excitation and inhibition and thus a decreased conductance (Rubin et al., 2015).

### Relationship to motion integration in MT

In MT, the integration of different motion directions has frequently been probed with the plaid stimuli (Movshon et al., 1985; Smith et al., 2005), comprised of superimposed gratings moving in different directions. Previous work has distinguished between pattern cells, which respond to the plaid motion direction, and component cells, which respond to the individual grating motion directions (Movshon et al., 1985).

In the terminology used here, a plaid stimulus moving in a neuron’s preferred direction entails component motion confined to the directional surround. Thus for a high-contrast plaid, the component gratings should suppress the neuron’s response, and this could contribute to the observed responses of component neurons. Furthermore, component-selective neurons have small direction centers (i.e. narrow tuning width), so that they do not integrate input from two gratings moving in very different directions (Rust et al., 2006; Tsui et al., 2010; Khawaja et al., 2013).

Pattern cells have broader direction tuning than component cells (Rust et al., 2006; Khawaja et al., 2013). Direction tuning, measured from the responses to individual motion directions, corresponds to the “minimal response field” in visual space, the region in which small stimuli can activate the cell; this measure does not change with contrast (Song and Li, 2008). Our measure of motion integration is not correlated with direction tuning width (Fig. 5E), and is best related to the “summation field size” in visual space, the size of a stimulus that best drives a cell before further size increases cause surround suppression. The summation field size, like our measure of motion integration, shrinks with contrast (Sceniak et al., 1999). We found a weak correlation between our motion integration index and the pattern index, which quantifies integration of plaid stimuli (Fig. 5D). These results suggest that the motion-domain summation field and pattern selectivity are linked, but that summation on its own is insufficient to account for pattern selectivity.

Pattern cells also show stronger suppression than component cells by stimuli moving opposite to their preferred directions (Rust et al., 2006). This suggests a direction-domain analogue of the “far surround” suppression that is found in the space domain; such suppression is also regulated by contrast both in the direction domain in MT (Pack et al., 2005) and in spatial surrounds in V1 (Schwabe et al., 2010). Our stimuli did not contain null-direction motion, and so they would not have probed this component of the MT receptive fields. Nevertheless, an inference from the existing data is that pattern cells in MT have both larger directional summation fields and larger (or stronger) directional surrounds.

It can be argued that random-dot stimuli are larger than gratings in the direction domain, as they activate a broader range of columns in V1 (Simoncelli and Heeger, 1998). Thus stimuli composed of multiple dots fields moving in different directions might elicit stronger suppression than grating stimuli containing a similar number of directions. Evidence in support of this idea comes from studies that use transparent motion stimuli, comprised of overlapping dot fields moving in two different directions. These stimuli evoke responses in MT that seem to reflect a suppression of responses to stimuli that straddle the preferred direction (Xiao and Huang, 2015), particularly for pattern cells (McDonald et al., 2014). One prediction of the current work is that such suppression should be weaker for low-contrast stimuli.

### Functional correlates of integration and suppression

A number of psychophysical studies have drawn a close link between contrast-dependent responses in MT and visual motion perception. For simple motion discrimination tasks, performance mirrors spatial processing in MT: for high-contrast stimuli, performance is worse for large than for small stimuli (Tadin et al., 2003; Liu et al., 2016). Similarly, motion perception can decrease at high contrasts when the stimulus speed is low, mirroring the contrast-dependent suppression found in MT (Pack et al., 2005; Seitz et al., 2008). In the direction domain, MT neurons exhibit higher null-direction suppression when the stimulus is high in contrast (Pack et al., 2005). This suggests further that suppressive influences are stronger for high-contrast stimuli, and there is some evidence that motion perception can worsen as the size of the stimulus increases in the direction domain (Treue et al., 2000; Dakin et al., 2005). Conversely, motion discrimination with noisy dots can sometimes improve at low contrast (Tadin et al., 2003). Our results predict the ability to integrate motion signals in the direction domain should systematically improve at low contrast, as has been found with manipulations of stimulus speed (Seitz et al., 2008) and spatial size (Tadin et al., 2003).

### Conclusion

A growing body of evidence points to a set of generic computations that are similar across brain regions (Creutzfeldt, 1977; Barlow, 1985; Miller, 2016) and across sensory modalities (Mountcastle, 1978; Pack and Bensmaia, 2015). Although this idea is attractive from a theoretical standpoint, it remains somewhat speculative. In this work, we have provided an experimental test of the genericity of one computational model by comparing results in MT with those obtained previously in V1. The qualitative pattern of results is similar, supporting the possibility that this model provides a more general framework for modulatory responses and integration in cortex.

## Acknowledgements

This work was supported by grants from the Canadian Institutes of Health Research to C.C.P. (PJT-148488) and L.D.L. (CGSD-121719), and NIH R01-EY11001 and the Gatsby Charitable Foundation (K.D.M.). We would like to thank Julie Coursol and the staff of the Animal Care Facility (Montreal Neurological Institute) for excellent technical support.

